# Temporal continuity of habitat strengthens covariation between plant and arbuscular mycorrhizal fungal communities

**DOI:** 10.1101/2024.07.03.601831

**Authors:** Ayako Shimono, Taiki Inoue, Hiroki Shiga, Kentaro Uchiyama, Yuki A. Yaida, Atushi Ushimaru, Tanaka Kenta

**Affiliations:** Department of Biology, Faculty of Science, Toho University, Chiba, Japan; Sugadaira Research Station, Mountain Science Center, University of Tsukuba, Nagano, Japan; Department of Forest Molecular Genetics and Biotechnology, Forestry and Forest Products Research Institute, Forest Research and Management Organization, Ibaraki, Japan; Graduate School of Human Development and Environment, Kobe University, Kobe, Japan

**Keywords:** arbuscular mycorrhizal fungi, covariation, semi-natural grassland, symbiosis

## Abstract

Old grassland is considered one of the most biodiverse terrestrial ecosystems. The habitat temporal continuity may promote the accumulation of host species and symbiont arbuscular mycorrhizal fungi (AMF) diversity and strengthen symbiont interactions. We tested the hypothesis that covariation between AMF and plant communities is stronger in old grasslands. We compared the relationship between AMF and plant communities in forests and in new and old semi-natural grasslands in Japan. DNA was extracted from bulk roots collected at each site and from roots of *Miscanthus sinensis* at each grassland site. AMF operational taxonomic unit was characterized on the basis of small subunit *rRNA* gene sequences. Old grasslands harboured the highest diversity of AMF among vegetation types. The AMF compositions were significantly related to those of plant species. The covariation of plant–AMF communities was stronger in old grasslands than in new grasslands. Individuals of *M. sinensis* were found to share AMF with bulk roots comprising different species. This suggests that AMF form a common mycorrhizal network among several coexisting plants of different species. Increasing AMF diversity in the shared rhizosphere has been proposed to lead to more efficient utilization of soil nutrients and to increase overall benefits of plant–AMF facilitation.

## Introduction

Mature grassland is one of the most biodiverse terrestrial ecosystems (Wilson *et al*., 2012). High plant species richness and the occurrence of habitat specialists—species that are characteristic of a habitat—are often associated with the long temporal continuity of grassland habitats (Inoue *et al*., 2021b; Noda *et al*., 2022). Old-growth grasslands are ecologically distinct from recently formed grasslands (Nerlekar & Veldman, 2020): old-growth grassland flora is distinct from early successional plant communities, even in managed grasslands. Once grasslands are destroyed or degraded, biodiversity recovery may require decades to centuries or may never occur (Nerlekar & Veldman, 2020; Buisson *et al*., 2022).

Mechanisms related to belowground mycorrhizal interactions are believed to control the diversity and composition of plant communities (van der Heijden *et al*., 2015; van der Putten *et al*., 2016; Tedersoo *et al*., 2020). Mycorrhizal fungi establish mutualistic associations with approximately 90% of vascular plant species (Brundrett & Tedersoo, 2018). Fungi receive carbohydrates from plants and provide plants with increased nutrient access and tolerance to abiotic and biotic stresses (Smith & Read, 2008). They also mediate plant interactions with other soil microbes, such as pathogens and mycorrhizosphere mutualists that protect against antagonists. Through these functions, mycorrhizal root symbionts influence belowground traits of plants and plant–plant interactions and regulate the diversity of plant communities (Tedersoo *et al*., 2020).

The most ubiquitous mycorrhizal symbiosis is with Arbuscular mycorrhizal fungi (AMF), which establish mutualistic associations with 70% to 80% of land plants, including grassland species (Brundrett & Tedersoo, 2018). Experimental evidence suggests that high AMF diversity enhances plant diversity and productivity as a result of more efficient exploitation of soil nutrients (van der Heijden *et al*., 1998; Wagg *et al*., 2011). There is some field evidence of positive local-scale correlation between plant and AMF diversities (Hiiesalu *et al*., 2014; Neuenkamp *et al*., 2018).

Although AMF typically have limited host specificity and nearly global distributions (Davison *et al*., 2015), many studies reported non-random associations between symbiotic AMF and host plant communities (Klironomos, 2003; Johnson *et al*., 2004; Koziol & Bever, 2015; Koziol & Bever, 2017) or AMF covariation with plant communities (Garcia de Leon *et al*., 2016; Neuenkamp *et al*., 2018). The driver–passenger hypothesis and the habitat hypothesis were proposed to explain the covariation or co-dependency between AMF and plant communities. The former states that plant communities drive AMF communities (or vice versa (Hart *et al*., 2001), and the latter states that the compositions of both AMF and plant communities are functions of environmental filtering (Zobel & Öpik, 2014). Kokkoris *et al*. (2020) suggested that covariation or co-dependency are likely to occur in homogeneous, stable environments where plants and AMF have co-occurred for a long time. Old-growth grasslands may serve as such stable environments. Numerous mechanisms are believed to control the AMF covariation or co-dependency with plant communities, but their respective roles and importance are still unclear.

Most of previous field studies that have evaluated the temporal covariation between AMF and plant communities assessed shifts in AMF communities along vegetation successional trajectories (Martinez-Garcia *et al*., 2015; Neuenkamp *et al*., 2018). They showed that the strength of plant–AMF correlation weakened during succession from grassland to forest following the cessation of grassland management, reflecting decreases in the proportion of obligate AM plants. However, little is known about the covariation between AMF and plant communities with temporal continuity of grassland (Garcia de Leon *et al*., 2016).

Here, we tested the hypothesis that the covariation between AMF and plants is strengthened in old grasslands, where plants and AMF have co-occurred for a long time. We compared the relationship between AMF and plant community compositions across forests and new and old semi-natural grasslands in Japan.

## Methods

### Study area and sites

Field surveys were conducted in ski run grasslands and adjacent secondary forests in the Sugadaira and Kirigamine highlands of central Japan, as in Inoue *et al*. (2021b, 2024). Using topographic maps from 1910–1912 and 1930–1937 and aerial photographs from 1946–1948, 1975, and 2010, we selected old and new grasslands and adjacent secondary forests as in Inoue *et al*. (2021b, 2024): old grasslands were those that had not been forested since 1910 and had persisted for at least 110 years, and new grasslands were those generated by clear-cutting of forests after 1910 to create ski runs and that had persisted for 45 to 89 years.

In Sugadaira, the mean annual temperature was 6.0 °C, ranging from 19.2 °C in July to −6.1 °C in January, and total precipitation was 1061 mm in 2017 (at Sugadaira Research Station, Mountain Science Center, University of Tsukuba, 1325 m a.s.l.). In Kirigamine, the mean annual temperature was 11.4 °C, ranging from 25.5 °C in July to −1.3 °C in January, and total precipitation was 1059 mm in 2017 (at a nearby JMA meteorological station, 36°02′N, 138°06′E, 760 m a.s.l.).

### Field investigation

In each study area, we selected seven old grasslands, six new grasslands, and six forests (Fig. 1; Table S1). The forest sites were situated adjacent to either old or new grassland sites. The forests were dominated by *Quercus* (Fagaceae), *Betula* (Betulaceae), or *Larix* (Pinaceae) trees. Grassland sites were located on ski runs, with the exception of one old grassland at the Sugadaira Research Station. A transect of 1 m × 20 m was set at each site, with at least 20 m from borders between vegetation types. In each transect, the presence of vascular plant species was recorded in summer (June to August) and autumn (September to October) in 2017 (Sugadaira) and 2018 (Kirigamine) by Inoue *et al*. (2021b, 2024). We used the data to examine the relationship between plant and AMF communities.

**Fig. 1.**
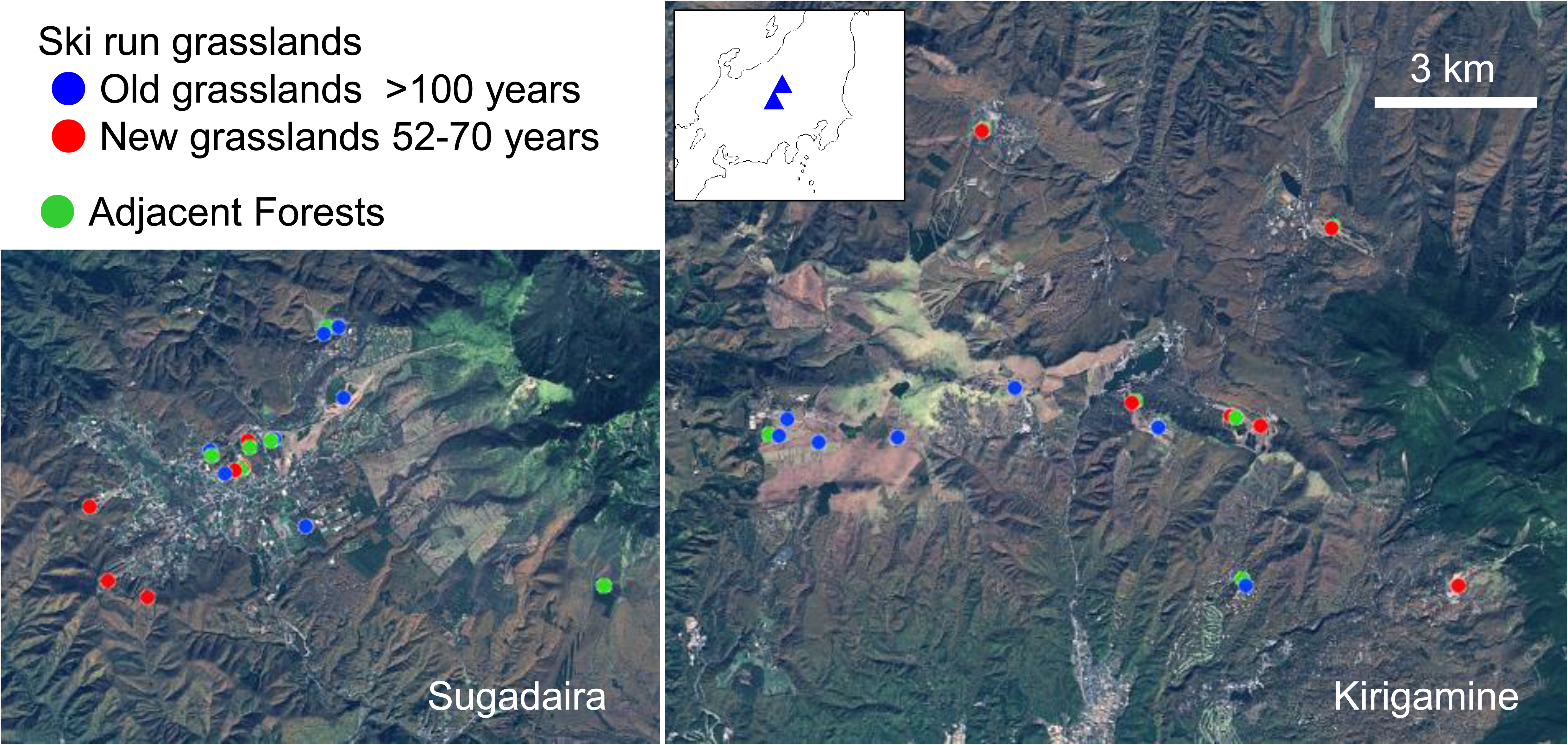
Locations of study sites.

We collected three soil cores (5 cm diameter, 10 cm deep) 5 m apart along each transect in September 2018 and combined them to form one composite sample. Bulk roots of unidentified plants were carefully separated within a few days after sampling. We also collected roots of three *Miscanthus sinensis* (Poaceae) plants near each transect, except in the forest sites, where this species was absent. This allowed us to compare AMF communities associated with the same host plants between the old and the new grasslands. All roots were thoroughly washed with water and stored in ethanol at 4 °C until extraction of DNA.

### Sequencing

DNA from each root sample was extracted with a DNeasy Plant Mini Kit (Qiagen, Hilden, Germany). The small subunit (SSU) *rRNA* gene was amplified with an AMF-specific primer pair (Sato *et al*., 2005), namely AMV4.5NF (5′-AAGCTCGTAGTTGAATTTCG-3′) and AMDGR (5′-CCCAACTATCCCTATTAATCAT-3′). Each primer was fused with a Fluidigm adapter: AMV4.5NF with CS1 (5′-ACACTGACGACATGGTTCTACA-3′) and AMDGR with CS2 (5′-TACGGTAGCAGAGACTTGGTCT-3′). PCR was performed in a total volume of 10 µL, containing 0.2 µM each primer, 0.2 mM each dNTP, 0.2 U KOD Plus Neo (Toyobo, Osaka, Japan), 1× KOD buffer, and ∼10 ng of template DNA. The reactions were run under a temperature profile of 94 °C for 2 min; 30 cycles of 98 °C for 10 s, 57 °C for 30 s, and 68 °C for 30 s; and a final extension at 68 °C for 3 min. Each amplicon was distinguished from others by a 10-bp molecular identifier, provided in the Fluidigm Access Array barcode library for Illumina sequencers 384 (single-direction) kit (Fluidigm, San Francisco, CA, USA). The second PCR was performed in a total volume of 10 µL, containing 0.2 µM of Fluidigm primer, 0.2 mM each dNTP, 0.2 U KOD Plus Neo, 1× KOD buffer, and 1 µL of PCR product. The reactions were run with a temperature profile of 94 °C for 2 min; 10 cycles of 98 °C for 10 s, 60 °C for 30 s, and 68 °C for 30 s; and a final extension at 68 °C for 3 min.

Each amplicon was purified with Agencourt AMPure XP (Beckman Coulter, Brea, CA, USA). All PCR products were then pooled to prepare a single product library. The library was quantified by qPCR using the Library Quantification Kit for Illumina sequencing platforms (KAPA Biosystems, Boston, MA, USA) on an Applied Biosystem 7500 real-time PCR system (Thermo Fisher Scientific, Waltham, MA, USA). MiSeq Standard (v. 3) 2 × 300 paired-end sequencing was performed on the Illumina MiSeq sequencer (Illumina, San Diego, CA, USA) with a 15% PhiX spike-in, following the manufacturer’s instructions. The sequenced files were deposited at NCBI Sequence Read Archive under the Bioproject accession PRJDB15351.

### Bioinformatics

The molecular ID and primer sequences were removed from the sequence reads in Claident v. 0.2 software (Tanabe & Toju, 2013). The forward and reverse sequence reads were fused, and then low-quality reads (≥10% of nucleotides had a quality of <30) and short reads (<200 bp) were eliminated. Chimeric reads were also eliminated in UCHIME v. 4.2.40 software (Edgar *et al*., 2011) with a minimum score of 0.1 to report a chimera. The sequence data were assembled in Assams v. 0.2 software (Tanabe & Toju, 2013) with a minimum similarity setting of 97%, and then consensus sequences were assigned as operational taxonomic units (OTUs). Since OTU sequences reconstructed from a small number of reads could be susceptible to sequencing errors, OTUs representing <10 reads were eliminated. A sample × OTU matrix, in which a cell represents the number of sequencing reads of an OTU in a sample, was obtained. Cells whose read counts represented <0.1% of the total read count of each OTU were removed to minimize effects of PCR and sequencing errors. To infer the taxonomy of the respective OTUs systematically, we prepared local BLAST databases from the “nt” database downloaded from the NCBI FTP server (http://www.ncbi.nlm.nih.gov/Ftp/) and conducted local BLAST searches in Claident (Tanabe & Toju, 2013). OTUs not assigned to the subphylum Glomeromycotina were discarded.

Each Glomeromycotina OTU was BLAST searched against SSU *rDNA* sequences in the Maarj*AM* database (Öpik *et al*., 2010). In the database, virtual taxa (VTXs) were defined for OTUs that were clustered at a 97% sequence similarity threshold for partial SSU *rDNA*. Each OTU was classified into a VTX as a match with a sequence similarity of ≥97% and an alignment length not differing from the length of the shorter of the query and subject (reference sequence) by >10%. When similarity values allowed the attribution to more than one VTX, the identification was based on the highest coverage value. When no match was found or for similarity <97%, an OTU was considered a putative new taxon. Some OTUs were classified into the same VTX, because the DNA region for OTUs at the variable site was shorter than that for the VTX, resulting in a finer division of the OTUs in this study than that of the VTXs in the Maarj*AM* database.

### Data analysis

From each sample we obtained 814 to 21 048 reads (average ± SD = 11 714 ± 4224; n = 116) and 23 to 187 OTUs (average ± SD = 112.6 ± 33.6; *n* = 116). To standardize the sample coverage, we conducted coverage-based rarefaction (Chao & Jost, 2012) using the *rareslope* and *rarefy* functions in the vegan package (Oksanen *et al*., 2020) in R v. 4.1.2 software. Coverage can be calculated from the slope of a traditional rarefaction curve. The sample coverage was set at 98.9%, because most of the samples with AMF could achieve this coverage. We discarded three samples (utg for bulk root and tug 1 and ctg 3 for *M. sinensis* roots) that did not achieve this coverage.

To examine the effects of region and habitat type on AMF or plant abundance, we conducted a GLM with a quasi-Poisson distribution and a log link. We used the number of OTUs of AMF or the number of plant species as the response variable and region (Sugadaira or Kirigamine) and habitat type (forest, new grassland, or old grassland) as the explanatory variables. Statistical analyses were conducted in R.

We calculated pairwise dissimilarity in OTU composition between all sites using the Bray–Curtis distance on rarefied AMF data and the Sørensen index on presence–absence plant data. Non-metric multidimensional scaling (NMDS) was used to visualize the separation of communities by the *monoMDS* function in the vegan package.

To determine whether the OTU composition of AMF communities and plant community compositions among habitat types or between regions differed statistically, we conducted a permutational analysis of variance (PERMANOVA) based on the dissimilarity index. Because spatial autocorrelations were detected, partial Mantel tests were conducted to separate the independent influences of grassland type and geographical distance on distance decay of community similarity. Geographical distances among sites were calculated as Euclidean distances. The partial Mantel test (with 10 000 permutations) was used to assess the influence of grassland type (new or old) on AMF or plant community similarities while holding geographical distance constant.

To assess covariation between plant species and AMF OTU compositions, we conducted two analyses. First, we fitted the first and second axis NMDS of the plant community on AMF NMDS ordinations with the *envfit* function in the vegan package with 10 000 permutations. Second, we performed Procrustes analysis (Peres-Neto & Jackson, 2001): the axes of the NMDS ordinations of AMF communities of bulk soil were rescaled and rotated to minimize the sum-of-squares deviations in relation to plant community NMDS axis scores. The residuals from the Procrustean solutions were used to assess differences in the strength of plant–AMF association between new and old grasslands.

The habitat indicator OTUs were identified by indicator species analyses that combined the relative abundance with the relative frequency of occurrence of an OTU in the vegetation types (Dufrene & Legendre, 1997), using the *indval* function in the labdsv package for R. Statistical significance was determined by 9999 randomization tests, and OTUs with *P* < 0.05 were considered significant indicator taxa of the habitat. When a corresponding VTX was found for an indicator OTU, the distribution of the VTX across ecosystem types was investigated in the Maarj*AM* database (Öpik *et al*., 2010). The registered sequences in each VTX were divided among forest, grassland, montane shrubland, anthropogenic, culture, and successional “Ecosystems” classifications and among the Asia, Europe, Africa, Oceania, North America, and South America “Continents” classifications in MaarjAM.

## Results

### Diversity of AMF

In total, 1 358 847 AMF reads were obtained and classified into 569 OTUs at a 97% sequence similarity threshold. After rarefaction, the number of AMF OTUs ranged from 5 to 106 across all but three samples (which did not achieve adequate coverage). The total number of vascular plant species was 367, ranging from 5 to 76 across all samples. The abundances of both AMF and plants were higher in grasslands than in forest (Fig. 2). Vegetation type had a significant effect on both AMF and plant abundances, but the effects of region and the interaction were not significant (Table 1). The number of AMF OTUs was positively correlated with the number of plant species (Fig. 3, *P* < 0.001), but only when forest sites were included.

**Fig. 2.**
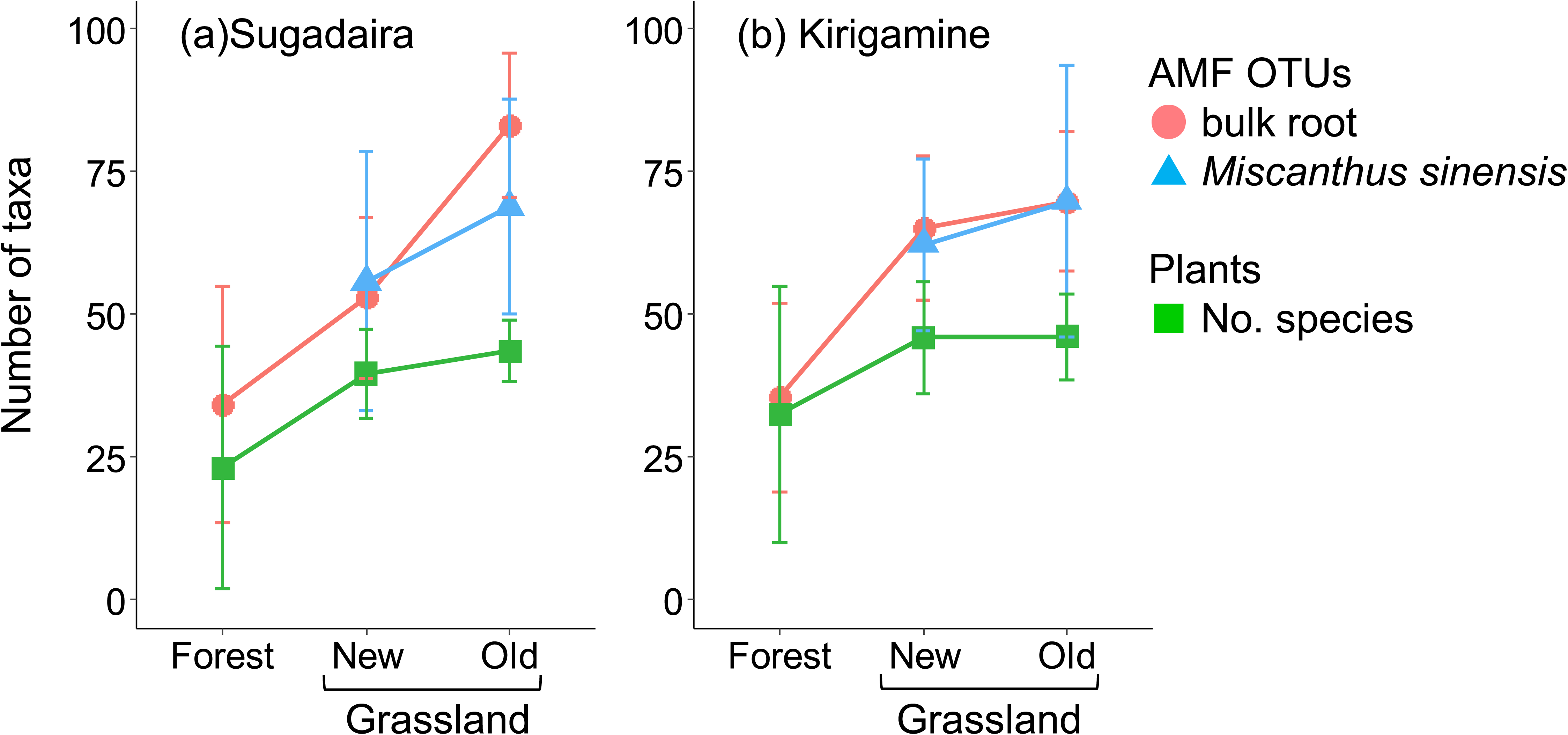
Mean numbers of arbuscular mycorrhizal fungal operational taxonomic units of bulk and *Miscanthus sinensis* roots and plant species from each habitat type in (a) Sugadaira and (b) Kirigamine. Bars indicate standard deviation.

**Fig. 3.**
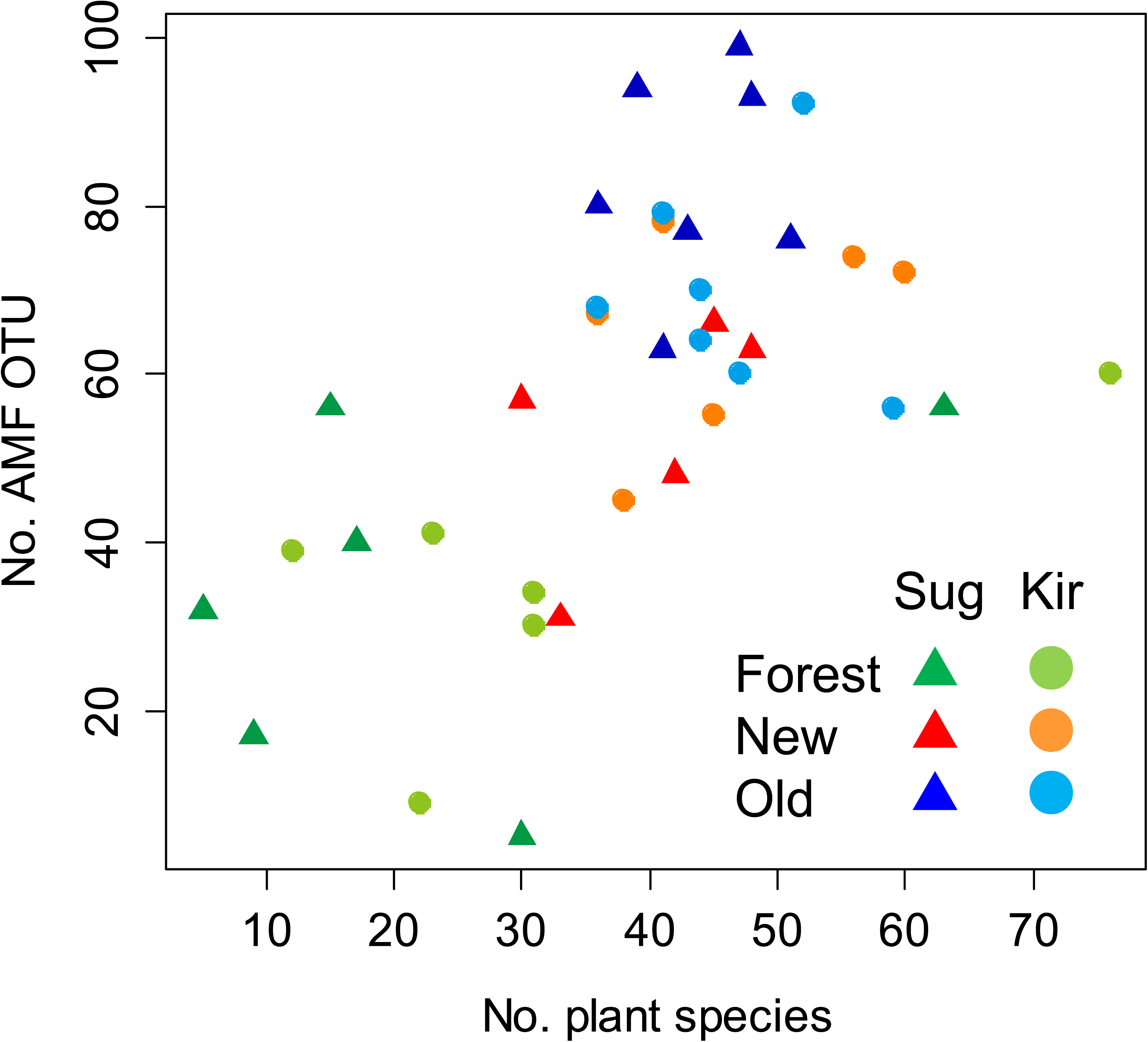
Relationship between number of plant species and arbuscular mycorrhizal fungal operational taxonomic units (OTUs) in Sugadaira and Kirigamine.

**Table 1.**
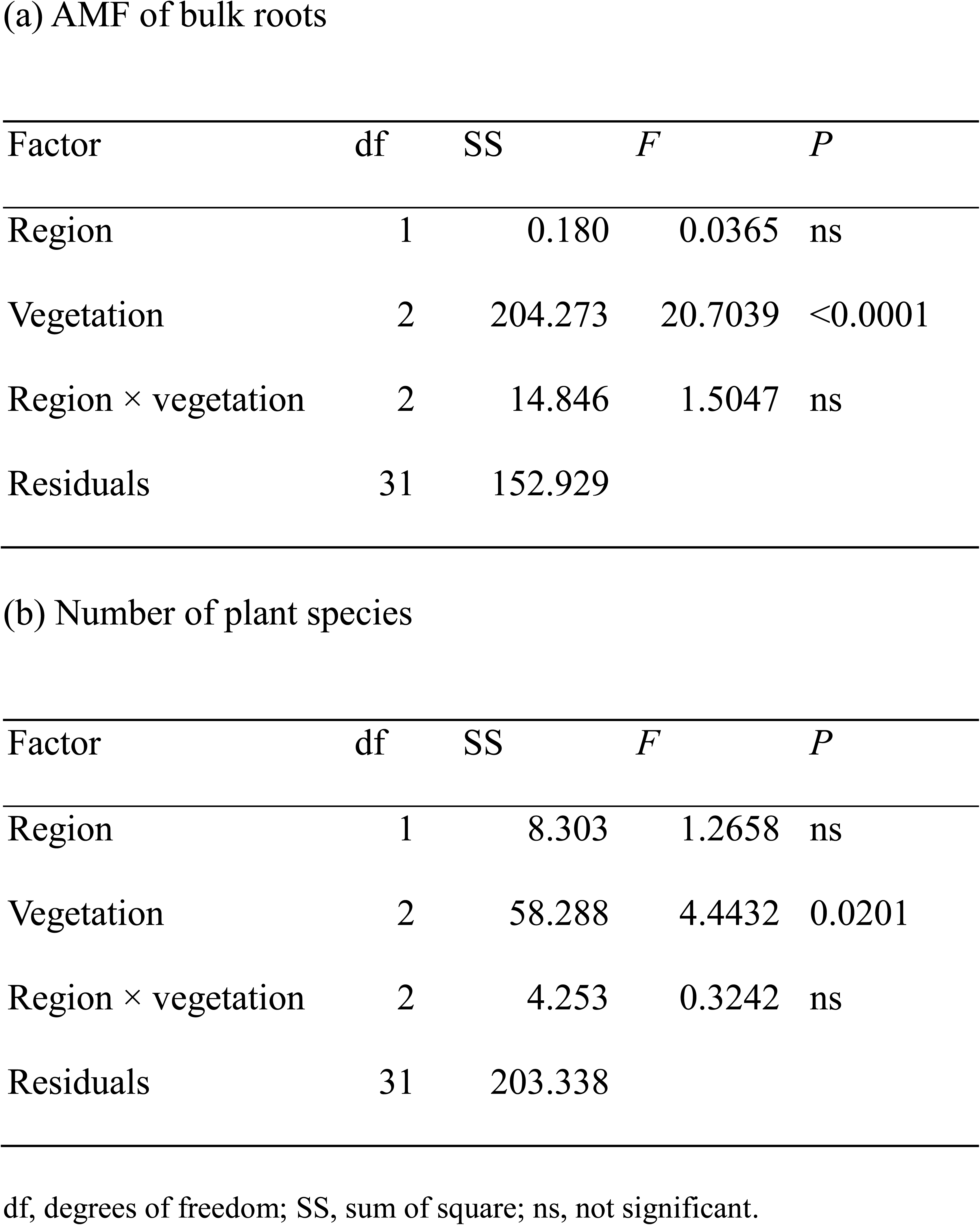
Effects of region (Sugadaira and Kirigamine) and vegetation (forest, new grassland and old grassland) on the number of operational taxonomic units (OTUs) of AMF of bulk roots (a) and number of plant species (b).

| Factor | df | SS | *F* | *P* |
| --- | --- | --- | --- | --- |
| Region | 1 | 0.180 | 0.0365 | ns |
| Vegetation | 2 | 204.273 | 20.7039 | <0.0001 |
| Region × vegetation | 2 | 14.846 | 1.5047 | ns |
| Residuals | 31 | 152.929 |  |  |

(b) Number of plant species
| Factor | df | SS | *F* | *P* |
| --- | --- | --- | --- | --- |
| Region | 1 | 8.303 | 1.2658 | ns |
| Vegetation | 2 | 58.288 | 4.4432 | 0.0201 |
| Region × vegetation | 2 | 4.253 | 0.3242 | ns |
| Residuals | 31 | 203.338 |  |  |

df, degrees of freedom; SS, sum of square; ns, not significant.

The numbers of AMF OTUs in both bulk and *M. sinensis* roots were significantly higher in old grasslands than in new grasslands (Fig. 2; Table 2). The number of AMF OTUs in *M. sinensis* roots was comparable to that in bulk roots. There were no significant effects of region, host, or their interaction on the number of AMF OTUs (Table 2).

**Table 2.** Effects of region (Sugadaira and Kirigamine), vegetation (new and old grasslands), and host (bulk and *Miscanthus sinensis*) on the number of AMF operational taxonomic units (OTUs) of bulk and *M. sinensis* roots in grassland habitats.

| Factor | df | SS | *F* | *P* |
| --- | --- | --- | --- | --- |
| Region | 1 | 1.74 | 0.3097 | ns |
| Vegetation | 1 | 54.59 | 9.7163 | 0.0024 |
| Host | 1 | 4.50 | 0.8010 | ns |
| Region × vegetation | 1 | 11.39 | 2.0276 | ns |
| Region × host | 1 | 2.34 | 0.4169 | ns |
| Vegetation × host | 1 | 1.84 | 0.3274 | ns |
| Region × vegetation × host | 1 | 5.99 | 1.0657 | ns |
| Residuals | 93 | 522.50 |  |  |

df, degrees of freedom; SS, sum of square; ns, not significant.

### Community structure and covariation

NMDS analysis revealed that the AMF OTUs of bulk roots and plant species compositions formed different clusters between forests and grasslands (Fig. 4). PERMANOVA indicated that both compositions differed significantly among habitats (both *P* < 0.001). Region had a marginal effect on AMF OTU composition (*P* = 0.07) and a significant effect on plant species composition (*P* < 0.05). The first NMDS axis of plant species composition was significantly correlated with the axes of AMF OTU composition (Fig. 4). PERMANOVA of the grassland habitats showed that grassland type had a marginal effect (*P* = 0.05) on the AMF OTU composition and a significant effect (*P* < 0.001) on the plant species composition (Fig. 5). Region had a significant effect on both AMF OTU (*P* < 0.05) and plant species (*P* < 0.001) compositions. The partial Mantel test confirmed that the AMF OTU and plant species compositions differed significantly between new and old grasslands (*P* = 0.02 for AMF, *P* < 0.001 for plants). The first and second NMDS axes of plant species composition were significantly correlated with those of AMF (Fig. 5).

**Fig. 4.**
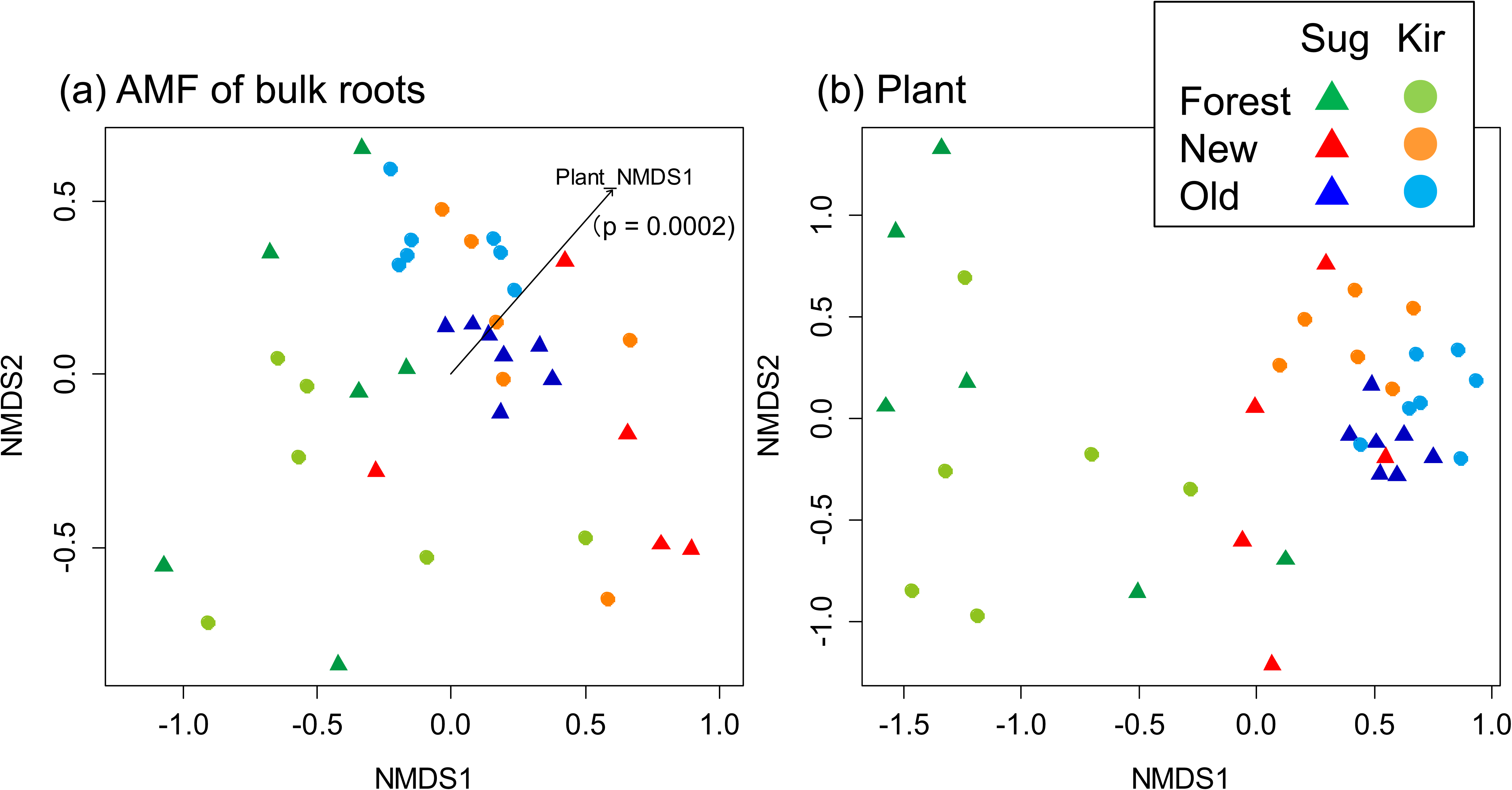
Non-parametric multidimensional scaling (NMDS) ordinations of (a) the arbuscular mycorrhizal fungal communities of bulk roots (Bray–Curtis dissimilarity index) and (b) plants (Sørensen dissimilarity index) from forests and old and new grasslands. Arrow marks the vector of the first axis of plant NMDS. *P*-values were obtained by 10 000 permutations.

**Fig. 5.**
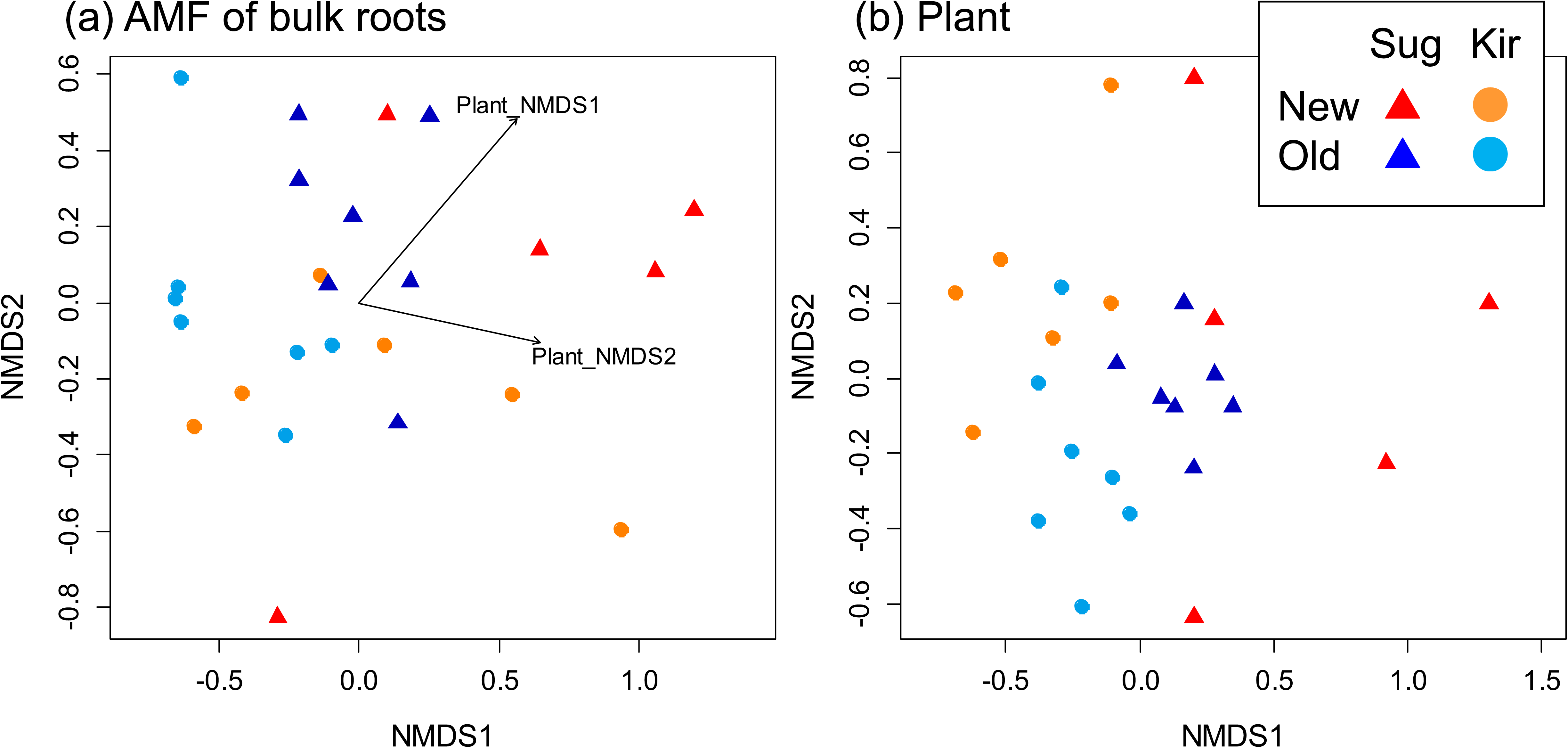
Non-parametric multidimensional scaling (NMDS) ordinations of (a) the arbuscular mycorrhizal fungal communities of bulk roots (Bray–Curtis dissimilarity index) and (b) plants (Sørensen dissimilarity index) from old and new grasslands. Arrows mark vectors of the first and second axes of plant NMDS. *P*-values were obtained by 10 000 permutations.

The AMF OTU composition of *M. sinensis* roots was similar that of bulk roots (Fig. 6). PERMANOVA showed that both grassland type and region significantly affected the AMF OTU composition of *M. sinensis* roots (both *P* < 0.001). The partial Mantel test confirmed that the AMF OTU composition of *M. sinensis* roots differed significantly between new and old grasslands (*P* < 0.001). The first and second NMDS axes of AMF OTUs of bulk roots and plant species compositions were significantly correlated with those of AMF of *M. sinensis* roots (Fig. 6).

**Fig. 6.**
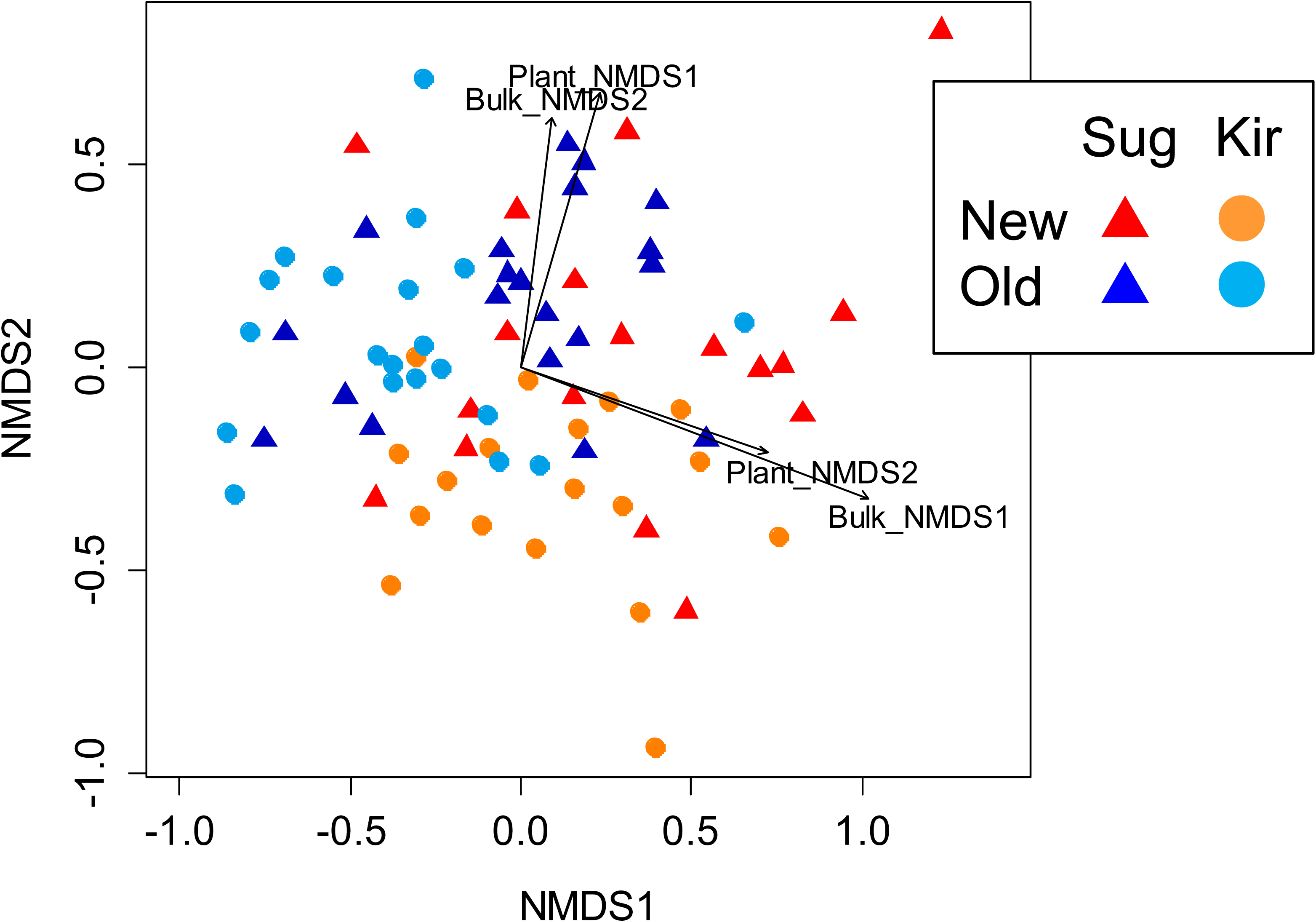
Non-parametric multidimensional scaling (NMDS) ordinations of the arbuscular mycorrhizal fungal communities of *Miscanthus sinensis* roots from old and new grasslands (Bray–Curtis dissimilarity index). Arrows mark vectors of the first and second axes of arbuscular mycorrhizal fungi of bulk root and plant NMDS. *P*-values were obtained by 10 000 permutations.

Procrustean randomization tests revealed highly significant covariation between plant species and AMF OTU compositions in bulk roots (Figs 7, S1). The Procrustean residuals between the AMF OTUs of bulk roots and plant species compositions were significantly smaller in the old grasslands than the new grasslands (*P* < 0.001; Fig. 7), suggesting that both communities were more strongly correlated in the old grasslands than in the new grasslands.

**Fig. 7.**
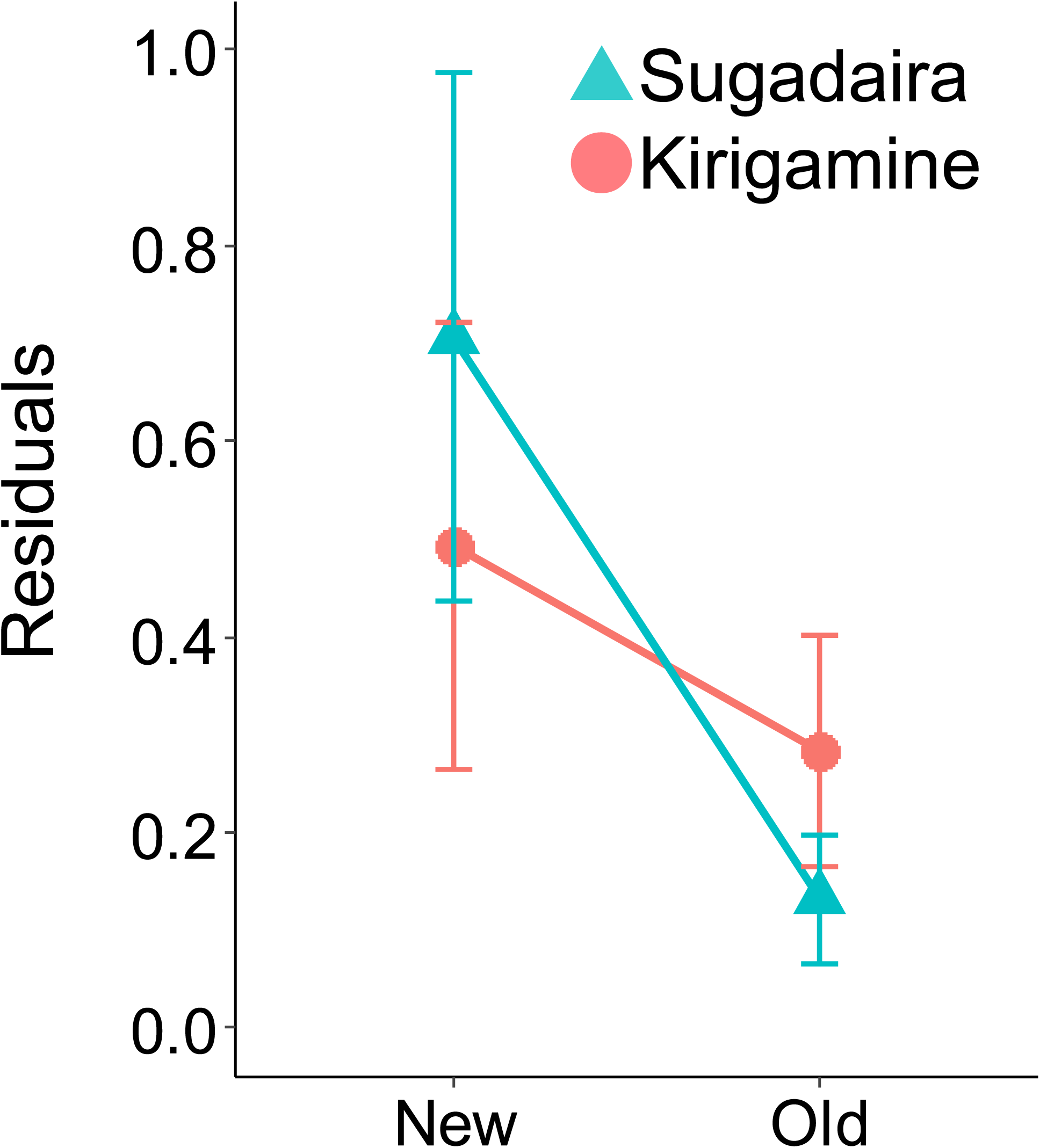
Procrustean residuals calculated by rotating the axes of a non-parametric multidimensional scaling (NMDS) ordination of arbuscular mycorrhizal fungal communities in bulk roots to minimize the sum-of-squares deviations from the axis scores of an NMDS ordination of plant communities. Bars indicate standard deviation.

### Indicator species analysis

From the analysis of bulk roots, we detected 3 OTUs as indicators of forest, 9 as indicators of new grasslands, and 12 as indicators of old grasslands (Table 3). Among these 24 OTUs, 19 corresponded to 14 VTXs in the Maarj*AM* database, where some OTUs were assigned to the same VTXs (Table 4). VTX074 and VTX080, detected as indicators of the forests here, were detected mostly in forests in Maarj*AM* (Table 4). VTXs other than VTX223 and VTX225, detected as indicators of new grasslands, were detected more in grasslands than in other ecosystems in Maarj*AM* (Table 4). On the other hand, VTXs other than VTX196 and VTX258, which were indicator taxa of old grasslands, were detected more in forests than in other ecosystems in Maarj*AM* (Table 4). VTX258 was detected only once in grassland in Japan. Other than this, no VTXs were specific to Asia.

**Table 3.**
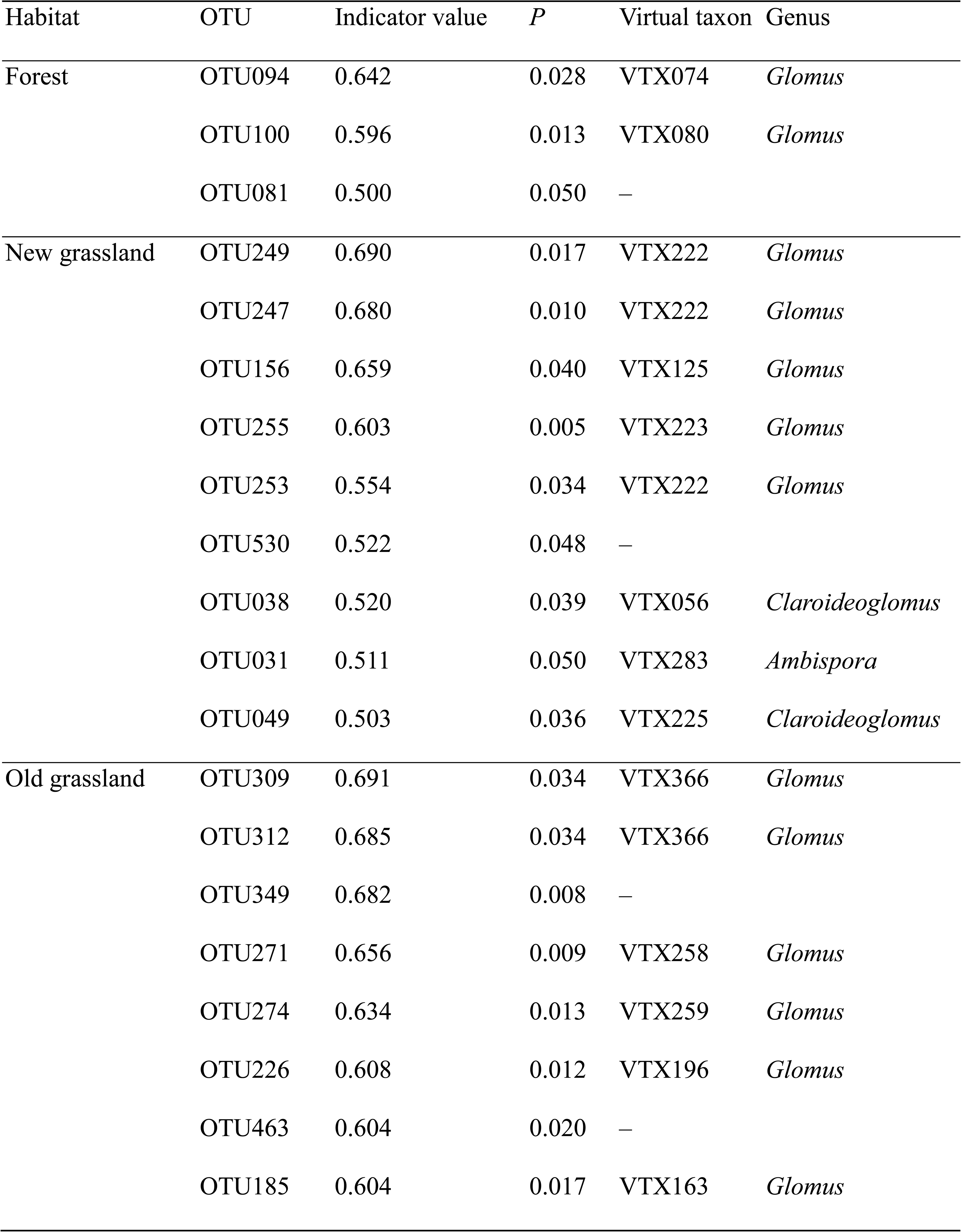
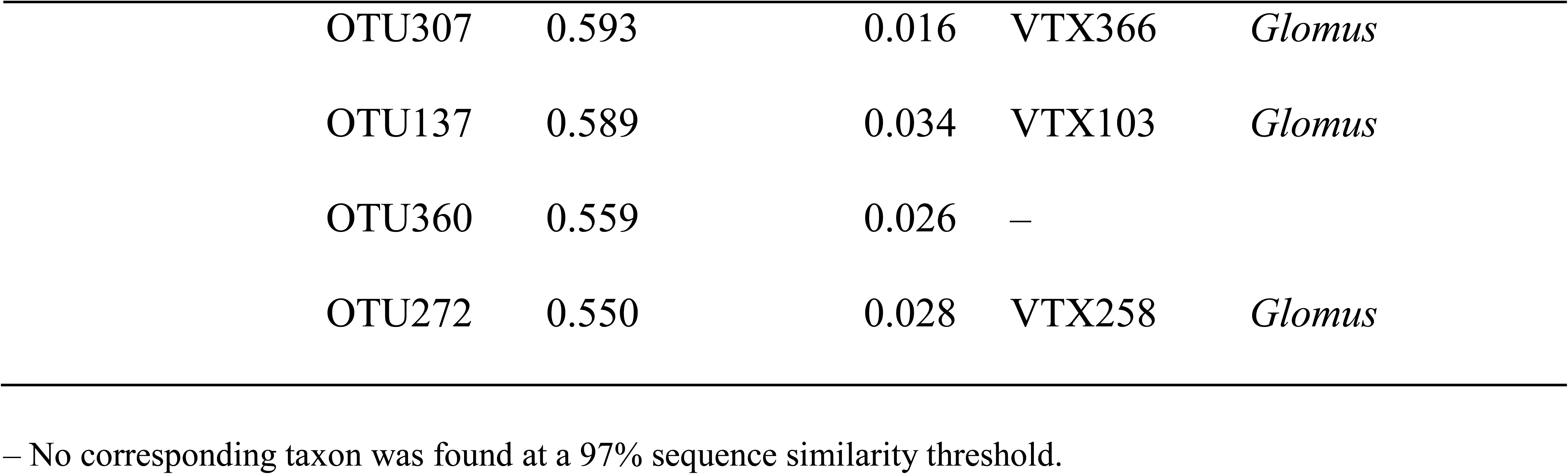
Indicator operational taxonomic units (OTUs) in each vegetation type and corresponding arbuscular mycorrhizal fungal virtual taxa (VTX) in Maarj*AM* database. Only species with significant indicator values are shown.

**Table 4.** Numbers of arbuscular mycorrhizal fungal virtual taxa (VTX) in Maarj*AM* database in seven ecosystems.

| Habitat | VTX | Forest | Grassland | Shrubland | Anthropogenic | Successional | Culture | Unspecified | Total |
| --- | --- | --- | --- | --- | --- | --- | --- | --- | --- |
| Forest | VTX074 | 243 | 82 | 57 | 16 | 21 |  |  | 419 |
|  | VTX080 | 122 | 26 | 6 | 2 | 4 |  | 3 | 163 |
|  | Total | 365 | 108 | 63 | 18 | 25 |  | 3 | 582 |
| New | VTX056 | 92 | 148 | 87 | 71 | 44 |  | 1 | 443 |
| grassland | VTX125 | 61 | 159 | 28 | 46 | 20 |  |  | 314 |
|  | VTX222 | 141 | 201 | 44 | 86 | 22 | 4 | 3 | 501 |
|  | VTX223 | 18 | 10 | 2 | 2 | 6 |  | 1 | 39 |
|  | VTX225 | 5 | 8 |  | 29 | 9 |  | 1 | 52 |
|  | VTX283 | 25 | 52 | 5 | 34 | 17 | 2 | 1 | 136 |
|  | Total | 342 | 578 | 166 | 268 | 118 | 6 | 7 | 1485 |
| Old | VTX103 | 68 | 28 | 31 | 22 | 4 |  |  | 153 |
| grassland | VTX163 | 161 | 94 | 76 | 46 | 38 |  | 4 | 419 |
|  | VTX196 | 38 | 46 | 1 | 4 | 3 |  |  | 92 |
|  | VTX258 |  | 1 |  |  |  |  |  | 1 |
|  | VTX259 | 7 |  |  |  | 3 |  |  | 10 |
|  | VTX366 | 23 | 11 | 39 | 19 |  |  |  | 92 |
|  | Total | 297 | 180 | 147 | 91 | 48 |  | 4 | 767 |

## Discussion

### AMF diversity and community

The number of AMF OTUs was higher in grassland than in forest. The forest sites were dominated by members of the Fagaceae, Betulaceae, and Pinaceae, which are hosts to ectomycorrhizal (EcM) fungi. Although many AM plants grow in forests, the AMF diversities among them are relatively low in temperate and boreal forests where EcM trees are dominant (Gerz *et al*., 2016). EcM fungi and litter accumulation reduce mycorrhizal root colonization of neighbouring AM plants (Becklin *et al*., 2012). In forest ecosystems with diverse plant taxa in Japan’s temperate region, AMF communities were constituted primarily of limited AMF taxa (19 OTUs), where three OTUs were dominant (Miyake *et al*., 2020), one of which was VTX080, an indicator taxon of forest here.

The old grasslands harboured significantly higher AMF diversity than the new grasslands in both bulk roots and *M. sinensis* roots. Even after more than 50 years in the new grasslands, the AMF diversity has not increased to that of the old grasslands, although previous studies reported that the number of AMF OTUs recovered in grasslands restored 20 or 50 years ago to those in ancient grasslands (Garcia de Leon *et al*., 2016; Honnay *et al*., 2017). Here, AMF diversity was significantly higher than in those previous studies, revealing the effect of temporal continuity of habitats at a longer timescale. In addition, the old grasslands of Sugadaira have been maintained for more than 300 years (Inoue *et al*., 2021a) and those of Kirigamine were deduced to have been maintained by burning management for several thousand years on the basis of Andosol soil analysis (Togashi *et al*., 2018). Prescribed burns during the dormant season have little effect on hemicryptophytes and geophytes on account of the low heat conductivity of the soil and rapid combustion of plant residues. Although Pressler *et al*. (2019) described a decrease in fungal biomass after fires in grasslands, whether prescribed fires or wildfires, in cases where the plant community composition remains largely intact, the AMF flora may easily recover after prescribed burns (Kivlin *et al*., 2021).

Plant abundance was higher in grassland than in forest, but there was no significant difference between old and new grasslands, although the old grasslands tended to harbour higher abundance in the Sugadaira Highland. In the Kirigamine Highland, browsing by the expanding sika deer (*Cervus nippon*) populations has reduced plant species diversity in recent decades (Uchida *et al*., 2020). This effect may explain the lack of difference in plant species abundance between old and new grasslands there.

Plant species composition differed significantly between old and new grasslands. Old grasslands serve as refugia for grassland-dependent native and rare species in both regions, and the species frequently distributed in old grasslands are native perennial forbs with underground storage organs (Inoue *et al*., 2024). These characteristics coincide with those of old-growth grasslands species in previous studies (Nerlekar & Veldman, 2020). AMF may be particularly important for perennial forbs with underground storage organs. Previous studies have suggested that the thick and long-lived roots of plants stimulate AMF symbiosis by supporting more AMF colonization (Eissenstat *et al*., 2015; Liu *et al*., 2015). Old grasslands’ plant species were described as having poor colonization ability, producing seeds dispersed by gravity or by ants, forming limited seed banks, and relying on clonal growth or asexual reproduction (Nerlekar & Veldman, 2020). These traits limit or delay the establishment of old grassland species in new grasslands, but the dispersal limitation may not explain the high diversity of AMF taxa in the old grasslands. The indicator OTUs of the grasslands are not limited to distribution within grassland ecosystems and are found on multiple continents. Generally, most AMF are cosmopolitan, and the same species can be found on multiple continents (Davison *et al*., 2015; Davison *et al*., 2018). Such a broad distribution pattern suggests a highly effective long-distance dispersal strategy. However, VTX258 was found only in grasslands in Japan. Few sequences of some OTUs have been published in previous studies based on a 97% sequence similarity threshold. The distributions of some taxa may be specific to certain habitats or regions.

### Relationship between AMF and plant communities

The diversity and composition of AMF OTUs were consistent between bulk roots and *M. sinensis* roots, indicating that *M. sinensis* individuals shared the same AMF with other species. AMF generally exhibit low host specificity and form a common mycorrhizal network between coexisting plants from different species, genera, and even families (Walder *et al*., 2012; van der Heijden *et al*., 2015). Plant interconnections provide pathways for the transfer of nutrients (He *et al*., 2009; Li *et al*., 2022). Experimental studies have demonstrated that plant coexistence and productivity increase with increasing numbers of AMF species as a result of the cumulative positive effect of each individual AMF species (van der Heijden *et al*., 1998; Wagg *et al*., 2011). Increasing AMF diversity has been proposed to lead to more efficient utilization of soil nutrients in the system (van der Heijden *et al*., 1998). In addition, different plant species showed distinct growth responses to different taxa of AMF (Klironomos, 2003; Hoeksema *et al*., 2018). In this sense, plant–AMF facilitation may be stronger in plant communities that share diverse AMF in the rhizosphere.

The NMDS axes of AMF OTU compositions were significantly correlated with those of plant species, and the covariations between AMF OTU and plant species compositions were stronger in the old grasslands than in the new grasslands. These non-random associations may result from shared environmental filters not driven by an interaction among partners (the habitat hypothesis). However, studies have revealed the interdependence between plant and AMF communities, which play a role in regulating the assembly of both symbiotic partners. Generally, variations in fungal community composition are better explained by plant species than by environmental conditions and geographical locations (Kokkoris *et al*., 2020). The plant–AMF correlation was strong when the abundance of obligate AM plants was high and declined as the proportion of facultative AM plants increased (Neuenkamp *et al*., 2018). Although our study does not provide information on the classification of plant species as obligate or facultative AM plants, we can hypothesize that obligate AM plants would increase in habitats where plants and AMF have coexisted for a long time. The extent to which the covariations between AMF and plant communities are interdependent (i.e., causally determine each other) remains unclear. Verifying the co-dependency between plant and AMF may provide us with a complementing framework that allows high plant species richness and the occurrence of habitat specialists in grasslands maintained over the long term.

## Acknowledgments

The authors are grateful to the land owners of our study sites—general foundation Nirei Kai, Sugadaira Bokujo, Hatsunekan, Imai-kan and Jozan-kan; and management companies Sugadaira Pine Beak Ski, Sugadaira Ski House Co., Ltd., Oku Davos Snow Park and HARE Sugadaira-Kogen Snow Resort, Hakuba Cortina Snow Resort, Tsugaike Kogen Snow Resort, Blanche Takayama Ski Resort, Shirakabako Royal Hill, Tokyu Resort Town Tateshina, Pilatus Tateshina Snow Resort, and Kurumayama-kogen Ski Resort—for allowing our field surveys.

## Author contributions

Ayako Shimono wrote the main manuscript text and conducted AMF experiments. Ayako Shimono, Taiki Inoue, and Kenta Tanaka conceived and designed the study. Ayako Shimono, Taiki Inoue, Hiroki Shiga, Yuki A. Yaida, Atushi Ushimaru, and Tanaka Kenta conducted field surveys of plant communities and collected root samples. Kentaro Uchiyama sequenced AMF *rRNA*. All authors read and approved the final manuscript.

## Data availability

The datasets generated during the study are available from the corresponding author on reasonable request. The sequenced files can be accessed at NCBI Sequence Read Archive under the Bioproject accession PRJDB15351.

---

**Supplemental Figure 1.**
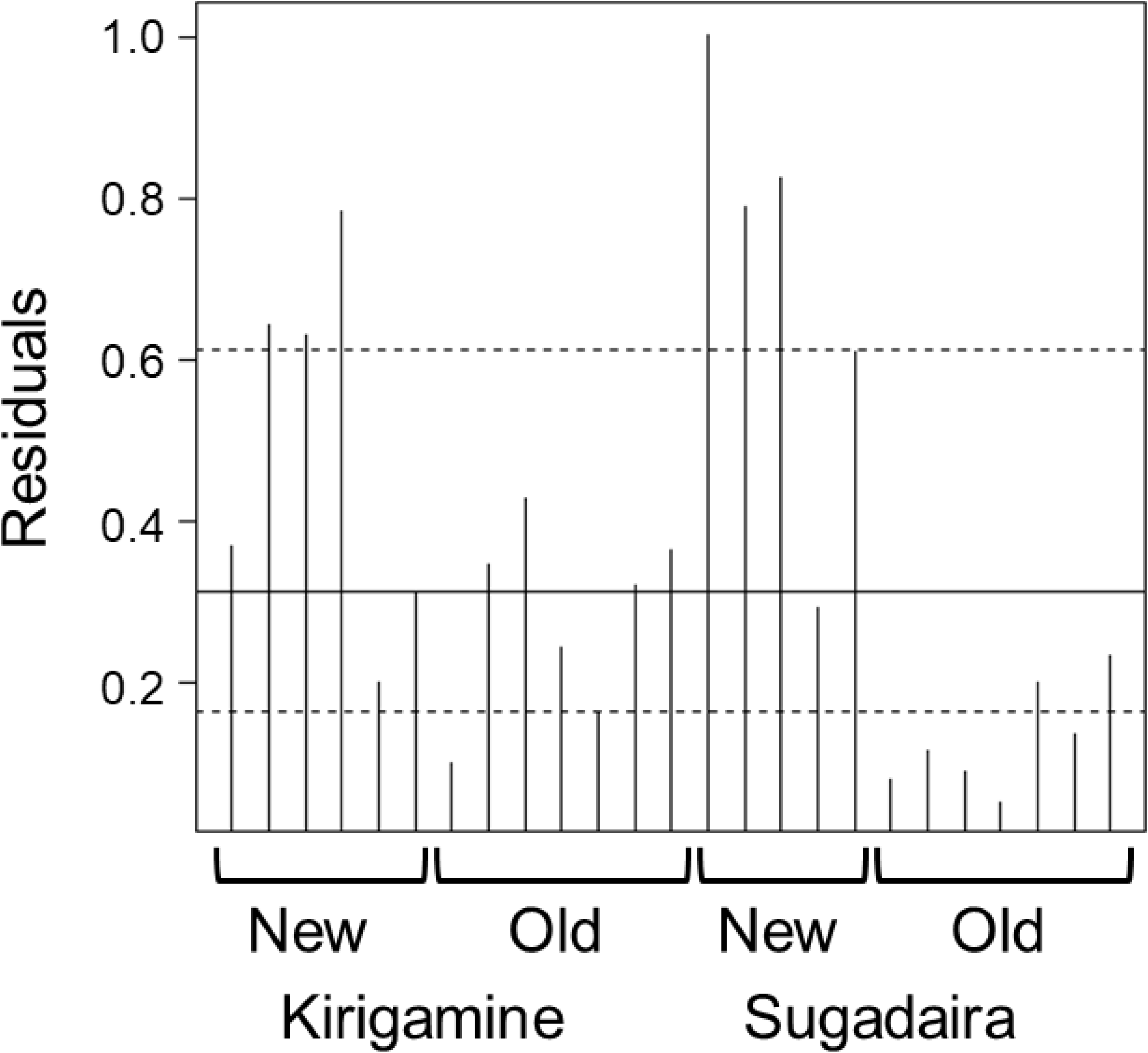
Histogram of the Procrustean residuals of each study site. Solid and dotted lines indicate median and interquartile range, respectively.

**Supplemental Table 1.**
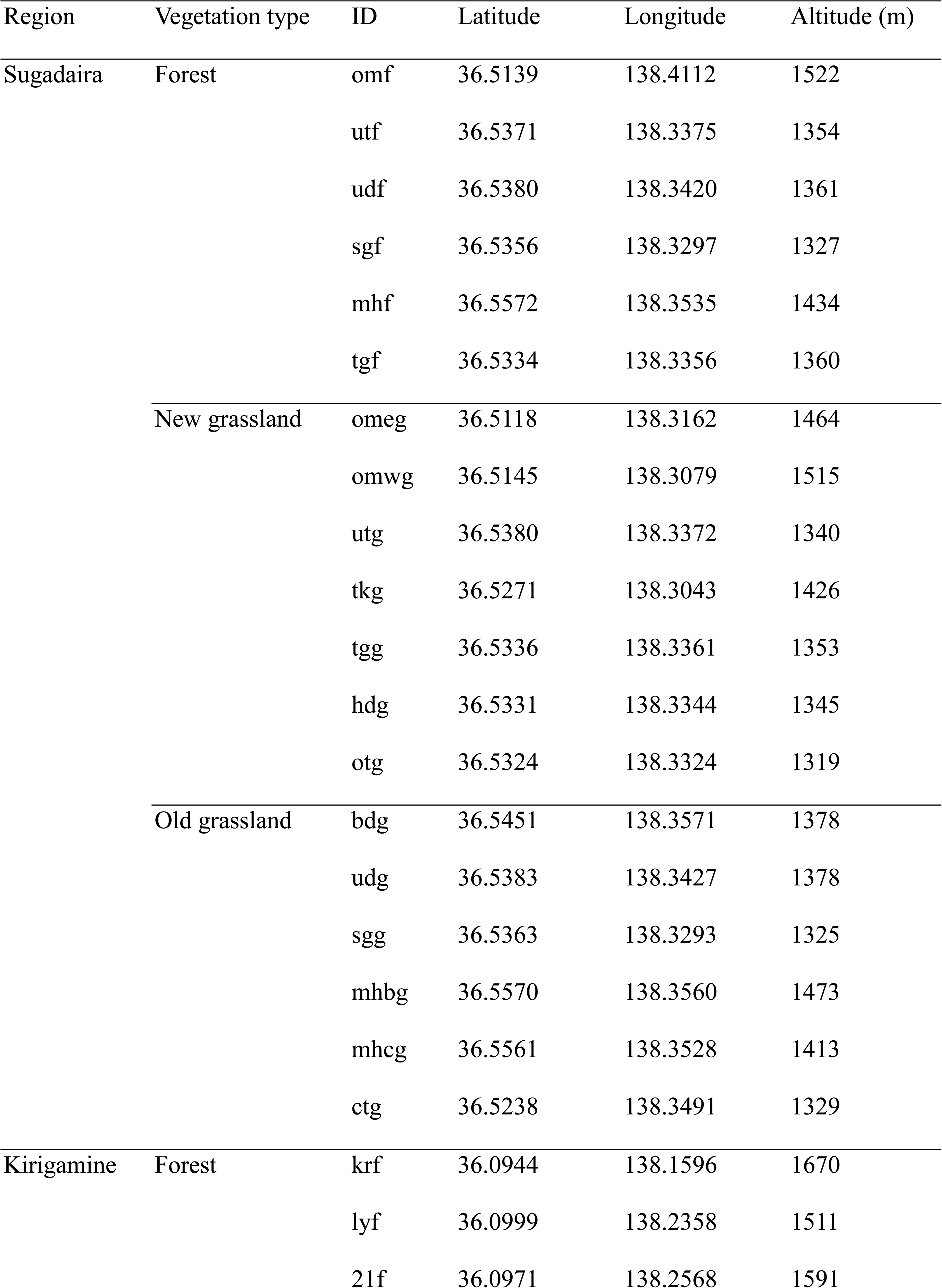
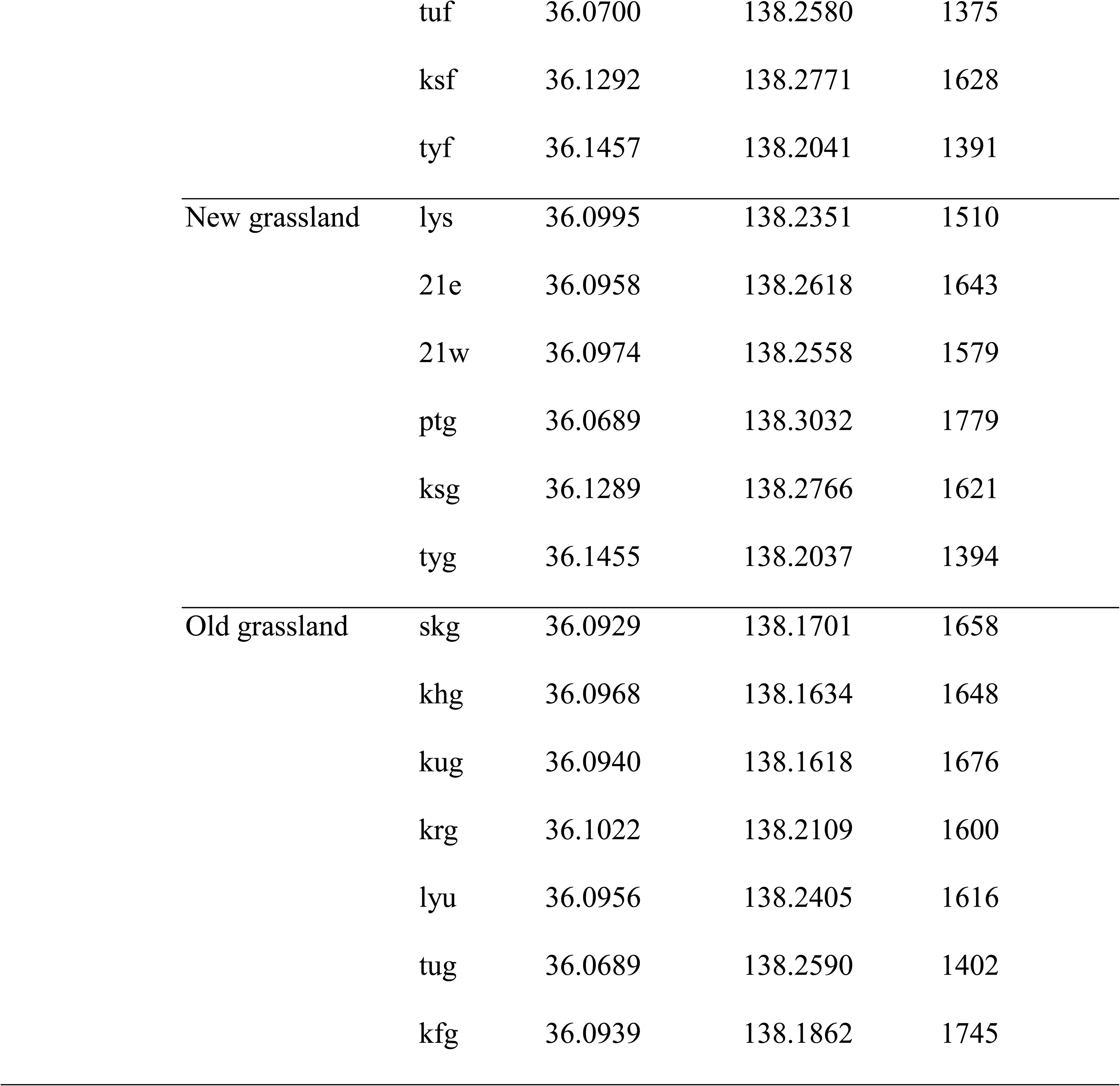
Locations and altitudes of the study sites in Sugadaira and Kirigamine.

## References

Becklin KM, Pallo ML, Galen C. 2012. Willows indirectly reduce arbuscular mycorrhizal fungal colonization in understorey communities. Journal of Ecology 100: 343–351.

Brundrett MC, Tedersoo L. 2018. Evolutionary history of mycorrhizal symbioses and global host plant diversity. New Phytologist 220: 1108–1115.

Buisson E, Archibald S, Fidelis A, Suding KN. 2022. Ancient grasslands guide ambitious goals in grassland restoration. Science 377: 594–598.

Chao A, Jost L. 2012. Coverage-based rarefaction and extrapolation: standardizing samples by completeness rather than size. Ecology 93: 2533–2547.

Davison J, Moora M, Öpik M, Ainsaar L, Ducousso M, Hiiesalu I, Jairus T, Johnson N, Jourand P, Kalamees R, et al. 2018. Microbial island biogeography: isolation shapes the life history characteristics but not diversity of root-symbiotic fungal communities. ISME Journal 12: 2211–2224.

Davison J, Moora M, Öpik M, Adholeya A, Ainsaar L, Ba A, Burla S, Diedhiou AG, Hiiesalu I, Jairus T, et al. 2015. Global assessment of arbuscular mycorrhizal fungus diversity reveals very low endemism. Science 349: 970–973.

Dufrene M, Legendre P. 1997. Species assemblages and indicator species: The need for a flexible asymmetrical approach. Ecological Monographs 67: 345–366.

Edgar RC, Haas BJ, Clemente JC, Quince C, Knight R. 2011. UCHIME improves sensitivity and speed of chimera detection. Bioinformatics 27: 2194–2200.

Eissenstat DM, Kucharski JM, Zadworny M, Adams TS, Koide RT. 2015. Linking root traits to nutrient foraging in arbuscular mycorrhizal trees in a temperate forest. New Phytologist 208: 114–124.

Garcia de Leon D, Moora M, Öpik M, Neuenkamp L, Gerz M, Jairus T, Vasar M, Bueno CG, Davison J, Zobel M. 2016. Symbiont dynamics during ecosystem succession: co-occurring plant and arbuscular mycorrhizal fungal communities. FEMS Microbiology Ecology 92.

Gerz M, Bueno CG, Zobel M, Moora M. 2016. Plant community mycorrhization in temperate forests and grasslands: relations with edaphic properties and plant diversity. Journal of Vegetation Science 27: 89–99.

Hart MM, Reader RJ, Klironomos JN. 2001. Life-history strategies of arbuscular mycorrhizal fungi in relation to their successional dynamics. Mycologia 93: 1186–1194.

He XH, Xu MG, Qiu GY, Zhou JB. 2009. Use of N-15 stable isotope to quantify nitrogen transfer between mycorrhizal plants. Journal of Plant Ecology 2: 107–118.

Hiiesalu I, Paertel M, Davison J, Gerhold P, Metsis M, Moora M, Öpik M, Vasar M, Zobel M, Wilson SD. 2014. Species richness of arbuscular mycorrhizal fungi: associations with grassland plant richness and biomass. New Phytologist 203: 233–244.

Hoeksema JD, Bever JD, Chakraborty S, Chaudhary VB, Gardes M, Gehring CA, Hart MM, Housworth EA, Kaonongbua W, Klironomos JN, et al. 2018. Evolutionary history of plant hosts and fungal symbionts predicts the strength of mycorrhizal mutualism. Communications Biology 1: 116.

Honnay O, Helsen K, Van Geel M. 2017. Plant community reassembly on restored semi-natural grasslands lags behind the assembly of the arbuscular mycorrhizal fungal communities. Biological Conservation 212: 196–208.

Inoue T, Okamoto T, Kenta T. 2021a. Qualitative and quantitative analyses of changes in the grassland area in the Sugadaira Highlands, central Japan: A case study of grassland decline in a national park. Japanese Journal of Conservation Ecology 26: 219–228.

Inoue T, Yaida YA, Uehara Y, Katsuhara KR, Kawai J, Takashima K, Miyamoto N, Yamamoto Y, Shimono A, Ushimaru A, et al. 2024. Effects of temporal grassland continuity on plant diversity and species composition in geographically diverse communities. submitted.

Inoue T, Yaida YA, Uehara Y, Katsuhara KR, Kawai J, Takashima K, Ushimaru A, Kenta T. 2021b. The effects of temporal continuities of grasslands on the diversity and species composition of plants. Ecological Research 36: 24–31.

Johnson D, Vandenkoornhuyse PJ, Leake JR, Gilbert L, Booth RE, Grime JP, Young JPW, Read DJ. 2004. Plant communities affect arbuscular mycorrhizal fungal diversity and community composition in grassland microcosms. New Phytologist 161: 503–515.

Kivlin SN, Harpe VR, Turner JH, Moore JAM, Moorhead LC, Beals KK, Hubert MM, Papes M, Schweitzer JA. 2021. Arbuscular mycorrhizal fungal response to fire and urbanization in the Great Smoky Mountains National Park. Elementa: Science of the Anthropocene 9: 1.

Klironomos JN. 2003. Variation in plant response to native and exotic arbuscular mycorrhizal fungi. Ecology 84: 2292–2301.

Kokkoris V, Lekberg Y, Antunes PM, Fahey C, Fordyce JA, Kivlin SN, Hart MM. 2020. Codependency between plant and arbuscular mycorrhizal fungal communities: what is the evidence? New Phytologist 228: 828–838.

Koziol L, Bever JD. 2015. Mycorrhizal response trades off with plant growth rate and increases with plant successional status. Ecology 96: 1768–1774.

Koziol L, Bever JD. 2017. The missing link in grassland restoration: arbuscular mycorrhizal fungi inoculation increases plant diversity and accelerates succession. Journal of Applied Ecology 54: 1301–1309.

Li CJ, Li HG, Hoffland E, Zhang FS, Zhang JL, Kuyper TW. 2022. Common mycorrhizal networks asymmetrically improve chickpea N and P acquisition and cause overyielding by a millet/chickpea mixture. Plant and Soil 472: 279–293.

Liu BT, Li HB, Zhu BA, Koide RT, Eissenstat DM, Guo DL. 2015. Complementarity in nutrient foraging strategies of absorptive fine roots and arbuscular mycorrhizal fungi across 14 coexisting subtropical tree species. New Phytologist 208: 125–136.

Martinez-Garcia LB, Richardson SJ, Tylianakis JM, Peltzer DA, Dickie IA. 2015. Host identity is a dominant driver of mycorrhizal fungal community composition during ecosystem development. New Phytologist 205: 1565–1576.

Miyake H, Ishitsuka S, Taniguchi T, Yamato M. 2020. Communities of arbuscular mycorrhizal fungi in forest ecosystems in Japan’s temperate region may be primarily constituted by limited fungal taxa. Mycorrhiza 30: 257–268.

Nerlekar AN, Veldman JW. 2020. High plant diversity and slow assembly of old-growth grasslands. Proceedings of the National Academy of Sciences of the United States of America 117: 18550–18556.

Neuenkamp L, Moora M, Öpik M, Davison J, Gerz M, Mannisto M, Jairus T, Vasar M, Zobel M. 2018. The role of plant mycorrhizal type and status in modulating the relationship between plant and arbuscular mycorrhizal fungal communities. New Phytologist 220: 1236–1247.

Noda A, Yamanouchi T, Kobayashi K, Nishihiro J. 2022. Temporal continuity and adjacent land use exert different effects on richness of grassland specialists and alien plants in semi-natural grassland. Applied Vegetation Science 25.

Oksanen J, Blanachet FG, Friendly M, Kindt R, Legendre P, McGlinn D, Minchin PR, O’Hara RB, Simpson GL, Solymos P, et al. 2020. Package ‘vegan’ community ecology package version 2.5.7. https://cran.ism.ac.jp/web/packages/vegan/vegan.pdf.

Öpik M, Vanatoa A, Vanatoa E, Moora M, Davison J, Kalwij JM, Reier U, Zobel M. 2010. The online database MaarjAM reveals global and ecosystemic distribution patterns in arbuscular mycorrhizal fungi (Glomeromycota). New Phytologist 188: 223–241.

Peres-Neto PR, Jackson DA. 2001. How well do multivariate data sets match? The advantages of a Procrustean superimposition approach over the Mantel test. Oecologia 129: 169–178.

Pressler Y, Moore JC, Cotrufo MF. 2019. Belowground community responses to fire: meta-analysis reveals contrasting responses of soil microorganisms and mesofauna. Oikos 128: 309–327.

Sato K, Suyama Y, Saito M, Sugawara K. 2005. A new primer for discrimination of arbuscular mycorrhizal fungi with polymerase chain reaction-denature gradient gel electrophoresis. Grassland Science 51: 179–181.

Smith SE, Read DJ. 2008. *Mycorrhizal Symbiosis*. Amsterdam: Academic Press.

Tanabe AS, Toju H. 2013. Two new computational methods for universal DNA barcoding: a benchmark using barcode sequences of bacteria, archaea, animals, fungi, and land plants. Plos One 8: e76910.

Tedersoo L, Bahram M, Zobel M. 2020. How mycorrhizal associations drive plant population and community biology. Science 367: eaba1223.

Togashi H, Okamoto T, Suka T. 2018. ^14^C ages and C/N ratios of the black soil overlying the Kirigamine Kogen Heights, central Japan. Bulletin Nagano Environment Conservation Research Institute 14: 7–12 (in Japanese).

Uchida K, Koyama A, Ozeki M, Iwasaki T, Nakahama N, Suka T. 2020. Does the local conservation practice of cultural ecosystem services maintain plant diversity in semi-natural grasslands in Kirigamine Plateau, Japan? Biological Conservation 250.

van der Heijden MGA, Klironomos JN, Ursic M, Moutoglis P, Streitwolf-Engel R, Boller T, Wiemken A, Sanders IR. 1998. Mycorrhizal fungal diversity determines plant biodiversity, ecosystem variability and productivity. Nature 396: 69–72.

van der Heijden MGA, Martin FM, Selosse M-A, Sanders IR. 2015. Mycorrhizal ecology and evolution: the past, the present, and the future. New Phytologist 205: 1406–1423.

van der Putten WH, Bradford MA, Brinkman EP, van de Voorde TFJ, Veen GF. 2016. Where, when and how plant-soil feedback matters in a changing world. Functional Ecology 30: 1109–1121.

Wagg C, Jansa J, Schmid B, van der Heijden MGA. 2011. Belowground biodiversity effects of plant symbionts support aboveground productivity. Ecology Letters 14: 1001–1009.

Walder F, Niemann H, Natarajan M, Lehmann MF, Boller T, Wiemken A. 2012. Mycorrhizal Networks: Common Goods of Plants Shared under Unequal Terms of Trade. Plant Physiology 159: 789-+.

Wilson JB, Peet RK, Dengler J, Partel M. 2012. Plant species richness: the world records. Journal of Vegetation Science 23: 796–802.

Zobel M, Öpik M. 2014. Plant and arbuscular mycorrhizal fungal (AMF) communities - which drives which? Journal of Vegetation Science 25: 1133–1140.

